# Characterization of betacoronavirus HKU-1 and OC43 internal proteins using a prototypic coronavirus

**DOI:** 10.1101/2025.04.23.650176

**Authors:** Chaminda D. Gunawardene, Isha Pandey, Shruti Chatterjee, Yoatzin Penaflor-Tellez, Abby Odle, Adriana Messyasz, Ricardo Rajsbaum, Alan Sariol, Lok-Yin Roy Wong

## Abstract

Coronaviruses express a repertoire of accessory proteins for evading host immune responses. A small internal (I) accessory protein is expressed by the genus *Betacoronavirus*. Previous studies reported that the I proteins of SARS-CoV, MERS-CoV and SARS-CoV-2 inhibit type I interferon (IFN-I) expression through distinct mechanisms and have different roles in pathogenesis. Human coronaviruses (hCoV) HKU1 and OC43 are betacoronaviruses that also encode the I protein as an accessory protein. Although hCoV-HKU1 and hCoV-OC43 predominantly cause common cold in healthy adults, susceptible individuals infected with these viruses can develop severe disease. However, the virulence factors contributing to pathogenesis after infection with common cold CoVs (CCCoVs) have not been fully characterized. In particular, the functions of the hCoV-HKU1 and hCoV-OC43 I proteins have not been previously reported. The lack of robust reverse genetic systems, tissue culture and animal models limit the study of hCoV-HKU1 and hCoV-OC43 pathogenesis. Here, we examine the role of the hCoV-HKU1 and hCoV-OC43 I proteins in pathogenesis using a prototypic coronavirus. We introduce the I proteins of hCoV-HKU1 and hCoV-OC43 independently to a neurotropic strain of mouse hepatitis virus (J2.2). J2.2 infection is well characterized with clearly defined immune responses which allows the study of I proteins in the context of authentic coronavirus infection. We show that the I protein of hCoV-HKU1, but not that of hCoV-OC43, ameliorates MHV-J2.2 pathogenesis while the I protein of MERS-CoV exacerbates disease. The presence of the hCoV-HKU1 I protein decreases virus titers and cytokine expression while the I protein of MERS-CoV leads to increased immune cell infiltration and virus titers in mice after J2.2 infection. Moreover, the I proteins of hCoV-HKU1 and hCoV-OC43 show different patterns of subcellular localization. Overall, our findings suggest that the I protein of different betacoronaviruses play unique roles in pathogenesis.

**Author Summary:** Factors governing the differences in the pathogenicity between highly pathogenic human coronaviruses (SARS-CoV, MERS-CoV and SARSR-CoV-2) and seasonal coronaviruses (hCoV-HKU1 and hCoV-OC43) are not completely understood. These differences are at least in part contributed to by the accessory proteins encoded between these two groups of human coronaviruses. The use of a heterologous coronavirus infection model provides an isogenic background for the direct comparison of viral proteins encoded by various coronaviruses. In this study, we compare the role of one of the accessory proteins encoded by betacoronaviruses, the I protein, in mediating disease outcome using a prototypic coronavirus. We demonstrate that the I protein of the highly pathogenic MERS-CoV but not that of the seasonal coronaviruses HKU1 and OC43 contributes to enhanced disease in the context of MHV-J2.2 infection, highlighting that virus-specific functions of accessory proteins encoded by different hCoVs.

## Introduction

Nine coronaviruses have been reported to infect humans up to date [1–9]. Among these 9 human coronaviruses (hCoVs), 5 belong to the genus *Betacoronavirus*, namely hCoV-OC43, hCoV-HKU1, Severe Acute Respiratory Syndrome Coronavirus (SARS-CoV), Middle East Respiratory Syndrome Coronavirus (MERS-CoV) and SARS-CoV-2. SARS-CoV, MERS-CoV and SARS-CoV-2 are known to cause acute respiratory distress syndrome (ARDS) in humans and are known as highly pathogenic hCoVs [1,10,11]. hCoV-HKU1 and hCoV-OC43 infection in humans primarily result in mild self-limiting upper respiratory tract infection with common cold symptoms that resolve without major complications [12,13]. Although hCoV-HKU1 and hCoV-OC43 are considered common cold CoVs (CCCoVs), susceptible populations may also develop lower respiratory tract infection and pneumonia upon infection [3,14]. The mechanism of pathogenesis after infection with hCoV-HKU1 and hCoV-OC43 has not been fully understood.

Coronaviruses express a set of accessory proteins that are known to modulate host immune responses and contribute to pathogenesis [15–19]. Although accessory proteins are conserved among members within the same subgenus, coronaviruses often encode a set of diverse accessory proteins as compared to viruses of other subgenera. The three highly pathogenic hCoVs belong to the subgenus *Sarbecovirus* (SARS-CoV and SARS-CoV-2) and *Merbecovirus* (MERS-CoV) while CCCoVs hCoV-HKU1 and hCoV-OC43 are classified in the genus *Embecovirus*. These viruses express a diverse repertoire of accessory proteins which may contribute to disease severity upon infection. In addition, highly pathogenic hCoVs often encode more accessory proteins than CCCoVs [20]. This prompted the possibility that the difference in pathogenicity between highly pathogenic hCoVs and CCCoVs may be attributed in part to the accessory proteins encoded by distinct hCoVs. While betacoronaviruses encode different accessory proteins across genera, a small internal (I) accessory gene that overlaps with the nucleocapsid (N) gene at the 3’ end of the genome is encoded by betacoronaviruses of various subgenera [21]. Expression of the I protein is dependent on the subgenomic RNA (sgRNA) of N as no transcriptional regulatory sequence (TRS) specific to the I gene has been identified. Translation of the I protein is initiated by leaky ribosomal scanning of the N sgRNA at an alternative (+1) reading frame. Previous studies showed that the I proteins of SARS-CoV (protein 9b), MERS-CoV (protein 8b) and SARS-CoV-2 (protein 9b) antagonize interferon (IFN) induction through distinct mechanisms [18,22,23]. Furthermore, we previously showed that protein 8b of MERS-CoV and protein 9b of SARS-CoV-2 play opposite role in modulating pathogenesis in experimentally infected animals [19]. However, the functions of the I proteins of other betacoronaviruses including hCoV-HKU1 and hCoV-OC43 have not been previously described.

Since robust reverse genetic systems and tissue culture and animal models are not currently available for hCoV-HKU1, we dissect the functions of the I proteins in the context of a prototypic mouse hepatitis virus (MHV). MHV strain J2.2 is neurotropic and causes demyelinating encephalomyelitis. J2.2 infection has been well characterized with clearly defined immune responses [24–28]. Previous reports demonstrated the feasibility of this model for studying accessory proteins of highly pathogenic hCoVs including SARS-CoV and MERS-CoV [19,29,30]. In addition, the J2.2 model serves as a platform for comparing the role of I proteins of betacoronaviruses in an isogenic virus background due to their virus-specific roles reported previously [19,31]. In this study, we investigate the function of the I proteins of hCoV-HKU1 and hCoV-OC43 in relation to pathogenesis. We introduced the I proteins of hCoV-HKU1 and hCoV-OC43 into J2.2. In addition, we included J2.2 expressing MERS-CoV protein 8b generated previously for comparison [19]. Consistent with previous reports [19,30], introduction of protein 8b to J2.2 resulted in enhanced disease in infected mice. However, the presence of hCoV-HKU1 I protein attenuated J2.2 infection with improved survival in infected mice, while the hCoV-OC43 I protein did not contribute to significant changes in disease outcomes after J2.2 infection. Further analysis revealed that the presence of protein 8b promotes neutrophil infiltration and virus replication in the brain while the I protein of hCoV-HKU1 decreased virus replication in the brain. Infection of bone marrow-derived macrophages (BMDM) with MHV-J2.2 encoding the hCoV-HKU1 I protein results in decreased cytokine expression. Consistently, transcriptomic analysis indicates that the I protein of hCoV-HKU1 reduces gene expression related to inflammatory response, IFN signaling, antiviral response and macrophage activation. Moreover, the I proteins of hCoV-HKU1 and hCoV-OC43 exhibit different patterns of localization in infected cells, suggesting a possible mechanism for the distinct disease outcomes. Taken together, our results revealed the contrasting roles of betacoronavirus I proteins in contribution to pathogenesis using a heterologous CoV infection model.

## Results

### Generation and validation of J2.2 viruses expressing the I proteins of hCoV-HKU1 and hCoV-OC43

To characterize the role of hCoV-HKU1 and hCoV-OC43 I proteins, we replaced open reading frame (ORF) 4 of J2.2 with the coding sequences of the I proteins of hCoV-HKU1 or hCoV-OC43 independently to generate J2.2 expressing the I protein of hCoV-HKU1 (J2.2-HKU1.I) or hCoV-OC43 (J2.2-OC43.I), respectively. The absence of ORF4 is not known to affect virus replication or result in attenuation of virulence [32]. A V5 epitope tag was included at the C-terminal end of the I protein coding sequences for I protein detection. To control for the addition of the I protein sequences, we further replaced ORF4 with sequences identical to that of the I proteins of hCoV-HKU1 or hCoV-OC43 except with nonsense mutations that abolish the expression of the I proteins to generate J2.2-HKU1.I* or J2.2-OC43.I* respectively. Furthermore, we included J2.2 expressing MERS-CoV protein 8b (J2.2-8b) and the corresponding control virus (J2.2-8b*) generated as previously described [19] (**Figure 1A**). The detailed sequences and the positions of nonsense mutations introduced to the control viruses are shown in **Figure S1**.

**Figure 1.**
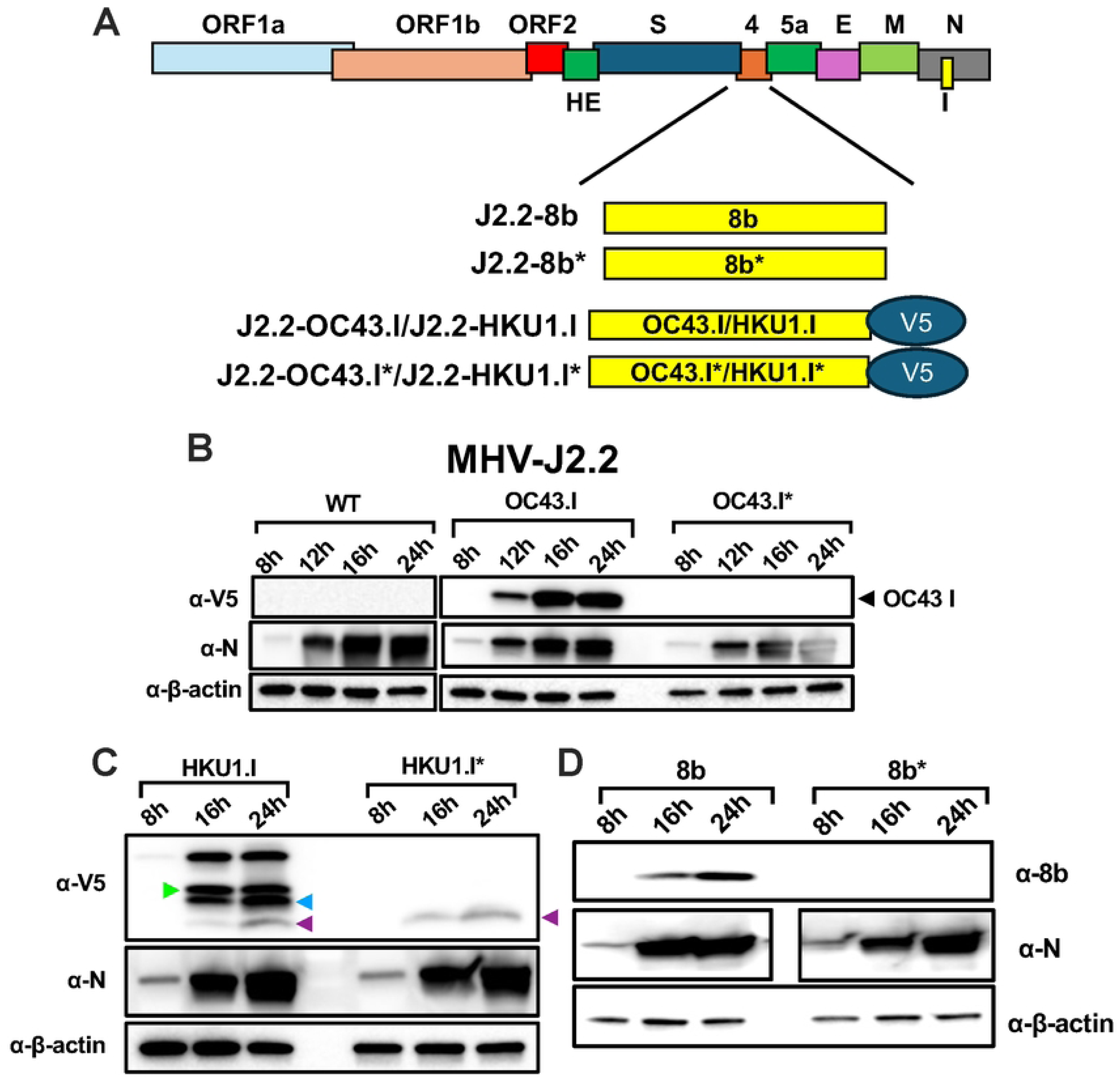
Verification of I protein expression in J2.2-infected cells. (A) Schematic diagram illustrating the introduction of I protein and the corresponding control sequences to replace ORF4 of J2.2. (B-D) 17Cl-1 cells were infected with the indicated viruses at a multiplicity of infection (MOI) of 0.01. Infected cells were harvested at the indicated time point. Cell lysates were subject to SDS-PAGE. The I proteins of hCoV-OC43 (B) and hCoV-HKU1 (C) were detected with an anti-V5 antibody (α-V5). Protein 8b (the I protein of MERS-CoV) was detected with an anti-8b antibody (α-8b) previously generated [18]. (D) Viral nucleocapsid protein (α-N) and β-actin (α-β-actin) were probed to control for virus replication and protein amount, respectively. The putative start codons for the three protein products (green, blue and purple arrowheads) are shown in **Figure S1**. Images are representative of three independent experiments.

We next assessed I protein expression in infected cells (**Figure 1B-D**). Cells infected with all J2.2 viruses exhibit increasing nucleocapsid (N) protein expression (α-N) from 8 hours post infection (hpi) to 24 hpi, indicating active virus replication. Wild-type (WT) J2.2-infected cells did not express any V5-tagged proteins (α-V5) (**Figure 1B, left panel**). J2.2-OC43.I-infected cells expressed the V5-tagged I protein as expected with increasing expression from 8-16 hpi and remained relatively constant from 16-24 hpi. Cells infected with J2.2-OC43.I* did not express any V5-tagged proteins (**Figure 1B, right panel**). Interestingly, cells infected with J2.2-HKU1.I expressed multiple V5-tagged proteins with increasing expression from 8-16 hpi and remain at peak expression till 24 hpi, suggesting that multiple proteins are expressed by the inserted hCoV-HKU1 *I* gene at start codons downstream of the first methionine residue in the context of J2.2 infection, which are detected by the C-terminal V5 tag. The top V5 band corresponds to the molecular weight of the full-length I protein of hCoV-HKU1. The putative start codons for the smaller bands are illustrated in **Figure S1**. V5-tagged proteins were not detected in J2.2-HKU1.I*-infected cells except for the one with the smallest molecular weight and lowest expression level (**Figure 1C, purple arrowhead**). The band intensity of this protein is much lower than other protein products, suggesting that a minor protein of small molecular weight is expressed during infection of J2.2-HKU1.I. Due to its low level of expression, we reason that this minor protein product would not complicate our analysis in a biologically relevant manner and that the phenotypic differences observed between J2.2-HKU1.I and J2.2-HKU1.I* can be attributed to other major protein products but not this minor protein. Consistent with previous reports [19], MERS-CoV protein 8b expression (α-8b) was detected in cells infected with J2.2-8b but not that infected with J2.2-8b* using an antibody targeting protein 8b that was raised previously [18] (**Figure 1D**).

### HKU1 I protein attenuates J2.2 infection *in vivo*

We next investigated if the I proteins play any role in regulating virus replication in a mouse fibroblast cell line 17Cl-1, which is reported to induce minimal interferon expression after MHV infection [33]. In general, J2.2 expressing the different I proteins did not contribute to significantly altered replication as compared to WT J2.2 and the corresponding control viruses except at 8 hpi, where J2.2-8b demonstrated a slight growth advantage (**Figure 2A**). To evaluate the function of the I proteins in relation to pathogenesis, we infected C57BL/6 mice with WT J2.2, J2.2-HKU1.I, J2.2-OC43.I and J2.2-8b. In agreement with previous reports, mice infected J2.2-8b developed more severe disease as compared to WT J2.2 [19,30]. However, J2.2-HKU1.I caused attenuated disease with reduced weight loss and clinical score and increased survival as compared to those infected with WT J2.2 (**Figure 2B**). To further confirm the impact of the I proteins on disease outcomes after J2.2 infection, we infected mice with J2.2 expressing the different I proteins and the corresponding control viruses. We again observed that mice infected with J2.2-HKU1.I developed attenuated disease as compared to those infected with J2.2-HKU1.I* (**Figure 3A**) while mice infected with J2.2-OC43.I and J2.2-OC43.I* developed similar clinical disease (**Figure 3B**). However, J2.2-8b caused significantly more severe disease with increased weight loss, mortality and clinical score as compared to those infected with J2.2-8b* (**Figure 3C**). These data indicate that the I protein of hCoV-HKU1 attenuates J2.2 infection while protein 8b of MERS-CoV exacerbates disease outcomes after infection, consistent with previous reports [19,30]. In addition, we observed that while most of the animals infected with J2.2-HKU1.I* and J2.2-OC43.I* succumbed to infection, all those infected with J2.2-8b* survived. Therefore, we compared the virulence of the three control viruses (J2.2-HKU1.I*, J2.2-OC43.I* and J2.2-8b*) to WT J2.2. J2.2-HKU1.I*, J2.2-OC43.I* and WT J2.2 caused similar disease in infected mice while J2.2-8b* is significantly attenuated as compared to WT J2.2 and the control viruses (**Figure S1**). These data underscore the importance of using viruses that control for the sequences being introduced for comparison, as introduction of sequences may have various effects on virulence.

**Figure 2.**
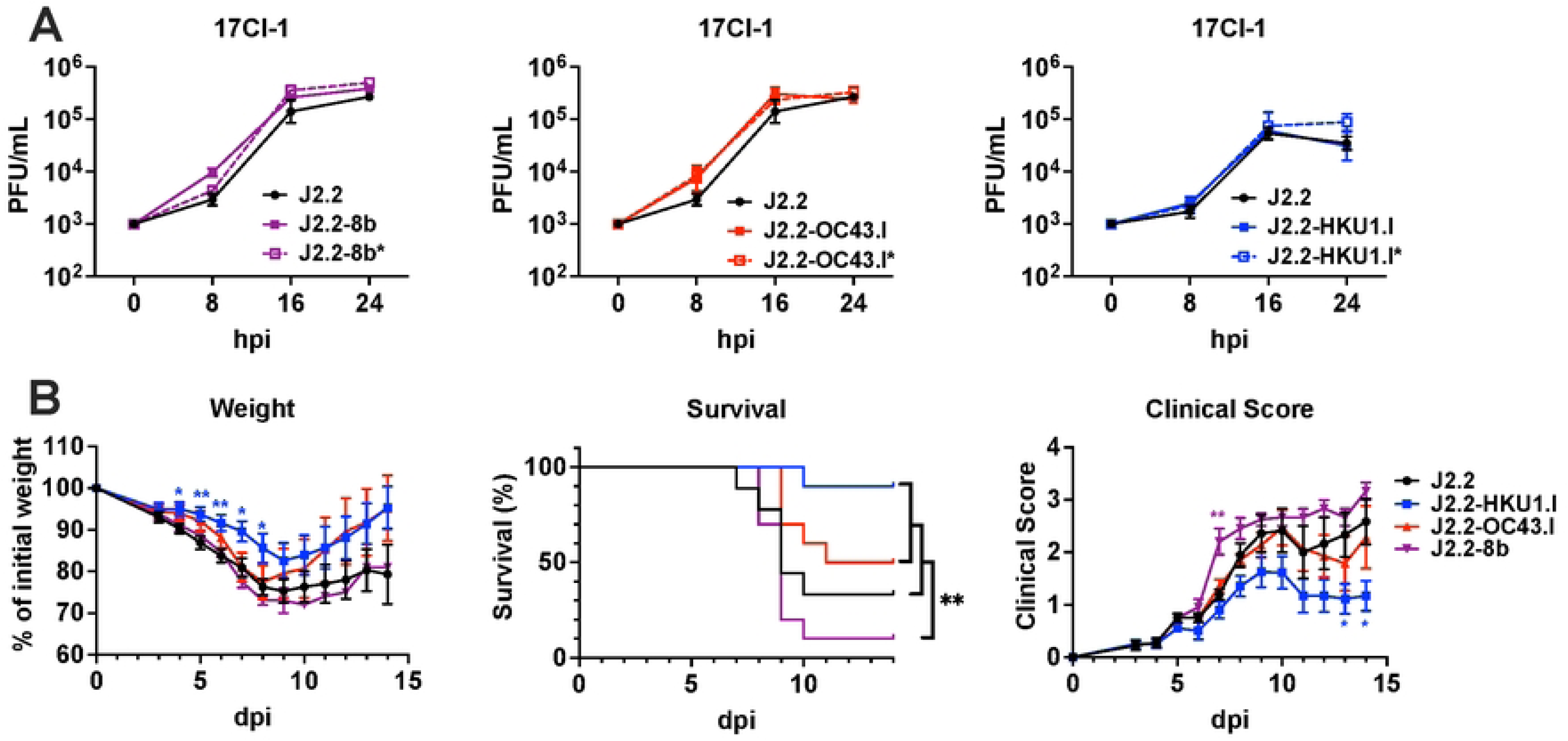
Effects of I protein on virus replication and disease outcomes. (A) 17Cl-1 cells were infected with WT J2.2 (J2.2, black solid line), J2.2 expressing the I protein of MERS-CoV (J2.2-8b, purple solid line) or J2.2 control virus for the I protein of MERS-CoV (J2.2-8b*, purple dashed line) (left panel); WT J2.2 (J2.2, black solid line), J2.2 expressing the I protein of hCoV-OC43 (J2.2-OC43.I, red solid line) or J2.2 control virus for the I protein of hCoV-OC43 (J2.2-OC43.I*, red dashed line) (middle panel); WT J2.2 (J2.2, black solid line), J2.2 expressing the I protein of hCoV-HKU1 (J2.2-HKU1.I, blue solid line) or J2.2 control virus for the I protein of hCoV-HKU1 (J2.2-HKU1.I*, blue dashed line) (right panel) at an MOI of 0.01. Cells and supernatant were collected at the indicated time points for plaque assay to determine virus titers (n = 3 for each data point). Data are representative of three independent experiments. Data points are shown as geometric mean ± geometric SD. (B) Percent of initial weight (left panel), survival (middle panel) and clinical scores (right panel) of C57BL/6 mice intracranially infected with 750 PFU of WT J2.2 (black), J2.2-HKU1.I (blue), J2.2-OC43.I (red) or J2.2-8b (purple). Data are pooled from at least two independent experiments (n ≥ 9 for each group). Data points are shown as mean ± SEM for the weight curve and the panel for clinical score. *P < 0.05, **P < 0.01 by Student’s t test at each time point as compared to J2.2. The color of asterisk represents the group of interest in the weight curve and the panel for clinical score. The P value in the survival curve was determined with log-rank (Mantel-Cox) test. **P<0.01.

**Figure 3.**
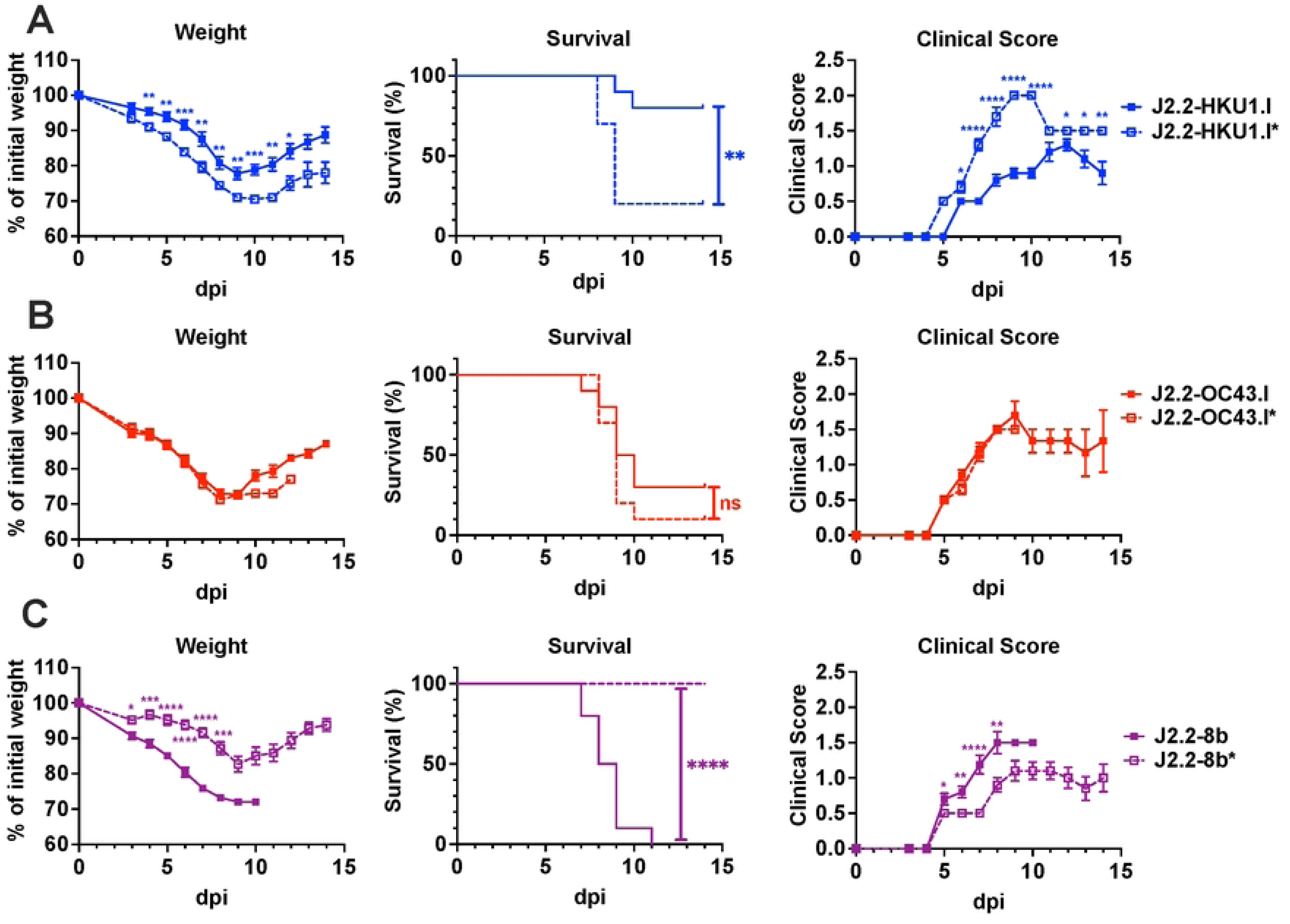
The I proteins of hCoV-HKU1, hCoV-OC43 and MERS-CoV play unique roles in J2.2 pathogenesis. Percent of initial weight (left panel), survival (middle panel) and clinical scores (right panel) of C57BL/6 mice intracranially infected with 750 PFU of J2.2-HKU1.I or J2.2-HKU1.I* (A); J2.2-OC43.I or J2.2-OC43.I* (B); J2.2-8b or J2.2-8b* (C). Data are pooled from at least two independent experiments (n = 10 for each group). Data points are shown as mean ± SEM for the weight curve and the panel for clinical score. *P < 0.05, **P < 0.01, ***P < 0.001, ****P < 0.0001 by Student’s t test at each time point. The P value in the survival curve was determined with log-rank (Mantel-Cox) test. **P<0.01, ****P<0.0001.

### hCoV-HKU1 I protein diminishes virus replication and dampens inflammatory responses in the brain

Based on these results, we determined the basis of attenuation afforded by the I protein of hCoV-HKU1. We assessed virus replication in the brain of mice infected with J2.2-HKU1.I or J2.2-HKU1.I*. Consistent with the observed attenuation, mice infected with J2.2-HKU1.I have significantly less virus in the brains as compared to those infected with J2.2-HKU1.I* at 3 and 7 days post infection (dpi) (**Figure 4A**). Similarly, we also identified significantly less viral genomic RNA (gRNA) in J2.2-HKU1.I-infected brains (**Figure 4C and D**). To characterize how the I protein of hCoV-HKU1 attenuates J2.2 infection, we infected BMDM isolated from C57BL/6 mice with J2.2-HKU1.I or J2.2-HKU1.I*. The levels of viral genomic RNA were not significantly different in BMDM infected with J2.2-HKU1.I or J2.2-HKU1.I* (**Figure 5A**). The expression levels of IL-6, IFNβ and ISG15 were significantly diminished in J2.2-HKU1.I-infected BMDM (**Figure 5B-E**), suggesting that the I protein of hCoV-HKU1 attenuates J2.2 infection by inducing less cytokine expression.

**Figure 4.**
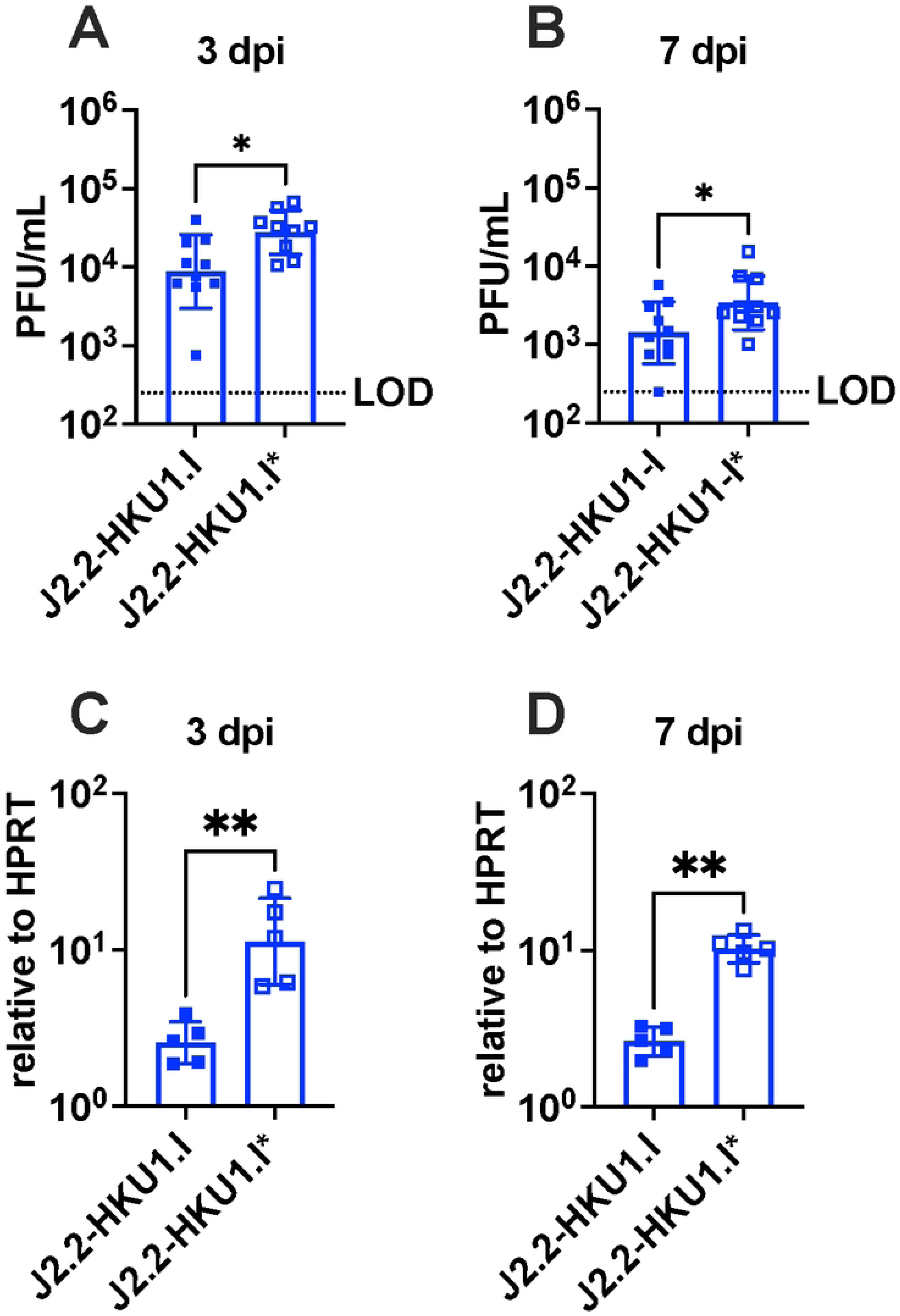
J2.2-HKU1.I replicates less efficiently in the brain. C57BL/6 mice were infected with 750 PFU of J2.2-HKU1.I or J2.2-HKU1.I* intracranially. The brains of infected mice were collected and homogenized at 3 (A, C) and 7 (B, D) dpi. (A, B) Infectious virus titers were determined by plaque assay in HeLa-MVR cells. (C, D) Expression levels of viral genomic RNA were assessed by qPCR. Data are pooled from two independent experiments in A and B and are representative of two independent experiments in C and D. Data points are shown as geometric mean ± geometric SD. Each point represents data obtained from an individual mouse. *P < 0.05, **P < 0.01 by Student’s t test.

**Figure 5.**
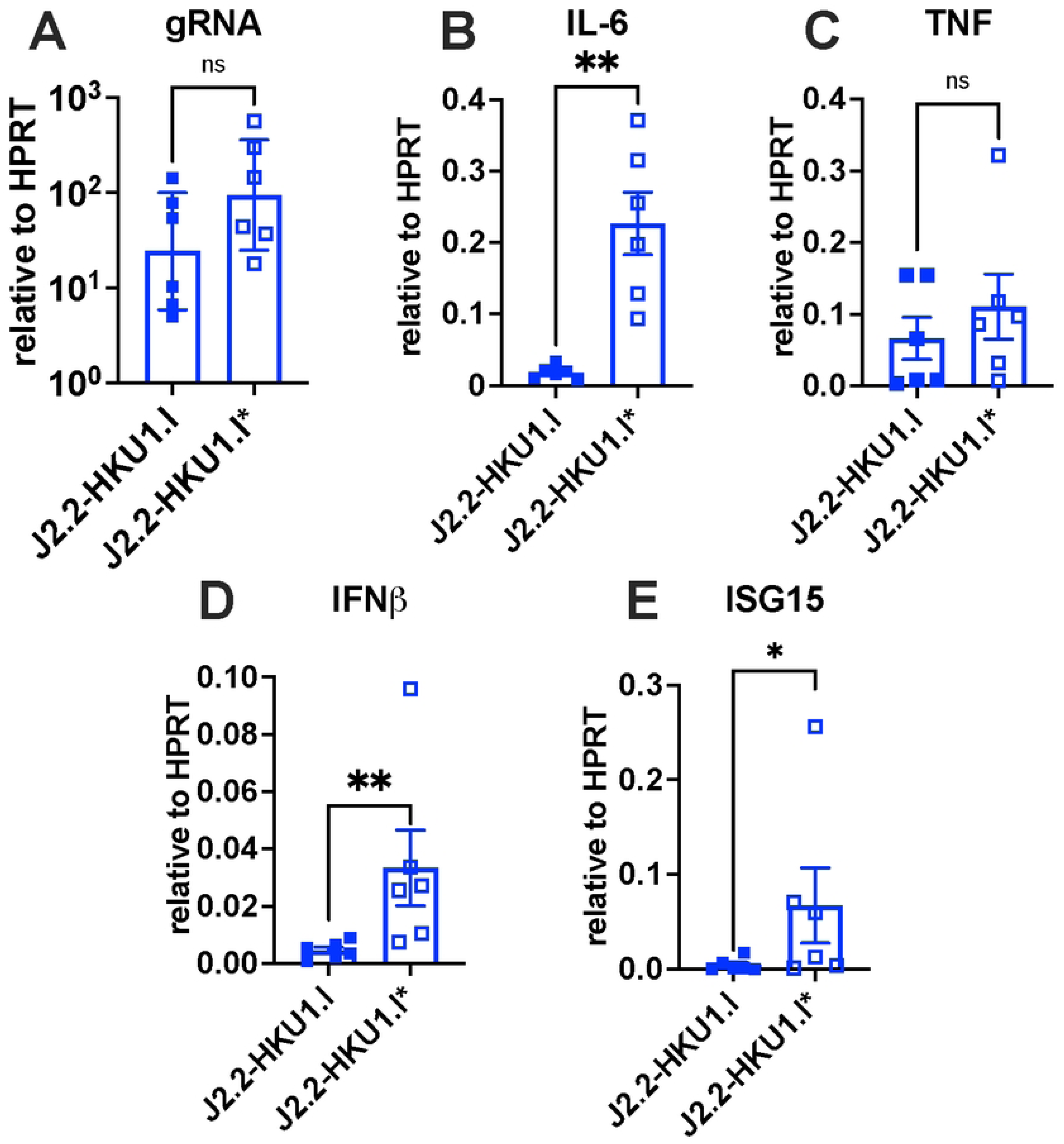
Decreased cytokine expression in J2.2-HKU1.I-infected BMDM. BMDM isolated from C57BL/6 mice were infected with J2.2-HKU1.I or J2.2-HKU1.I* at an MOI of 0.01. Infected BMDM were collected for RNA isolation and determination of cytokine expression by qPCR at 16 hpi. Data are pooled from two independent experiments. Each point represents data obtained from an individual biological replicate. (A) Data points are shown as geometric mean ± geometric SD. (B-E) Data points are shown as mean ± SEM. *P < 0.05, **P < 0.01 by Student’s t test.

To identify additional changes in host responses after J2.2-HKU1.I and J2.2-HKU1.I* infection, we performed RNA sequencing on J2.2-HKU1.I- and J2.2-HKU1.I*-infected brains. We observed 602 differentially expressed genes (523 upregulated and 79 downregulated; adjusted p values<0.05) in mice infected with J2.2-HKU1.I as compared to those infected with J2.2-HKU1.I* (**Figure S2A**). Genes related to cytokine storm, IFN signaling and neutrophil degranulation were significantly downregulated in J2.2-HKU1.I-infected mice (**Figure S2B-D**). Furthermore, we performed Ingenuity Pathway Analysis (IPA) to reveal significantly altered biological pathways (**Figure S2E**). In addition to the pathways shown in **Figure S2B-D**, J2.2-HKU1.I-infected mice express lower levels of genes involved in ISGylation signaling pathway, antiviral response and macrophage activation pathway. These data suggest that J2.2-HKU1.I-infected mice mount modest antiviral and inflammatory responses with attenuated disease as compared to those infected with J2.2-HKU1.I*.

### MERS-CoV protein 8b but not the hCoV-HKU1 I protein promotes virus replication and immune cell infiltration in the brain

We and others demonstrated previously that protein 8b enhances disease after J2.2 infection [19,30]. Here, we further characterize and compare the role of protein 8b and the hCoV-HKU1 I protein in modulating disease outcomes after J2.2 infection at the cellular level. We observed that the brains of mice infected with J2.2-8b harbors more infectious virus as compared to those infected with J2.2-8b* at 3 and 7 dpi (**Figure 6A**). This is in agreement with previous reports [19,30] and our observation (**Figure 3**) that protein 8b promotes severe disease with concomitant increase in virus replication in the brain in the context of J2.2 infection. Since J2.2 infection is known to recruit immune cells to the brain, we next evaluated the levels of different immune cell populations in the brains of mice infected with the I protein expressing viruses and their corresponding control viruses. Mice infected with J2.2-8b showed an increase in frequency and number of neutrophils at 3 dpi (**Figure 6A and 6B**), suggesting that the presence of protein 8b promotes the recruitment of pro-inflammatory neutrophils, possibly a consequence of increased virus replication, which contributes to severe disease. We also observed lower frequency and number of microglia at 7 dpi in the brains of J2.2-8b-infected mice as compared to those infected with J2.2-8b* (**Figure 6C and 6D**). This suggests that protein 8b may reduce microgliosis which leads to worse disease outcomes as microglia are important for protection during acute MHV infection and recovery after virus clearance [24,25]. In contrast, we did not identify significant changes in the frequency and number of neutrophils or microglia or other immune cells in the brains of mice infected with J2.2-HKU1.I compared to those infected with J2.2-HKU1.I* (**Figure 6 and S3**). These data provide a plausible explanation for the role of protein 8b but not the hCoV-HKU1 I protein in exacerbating J2.2 infection.

**Figure 6.**
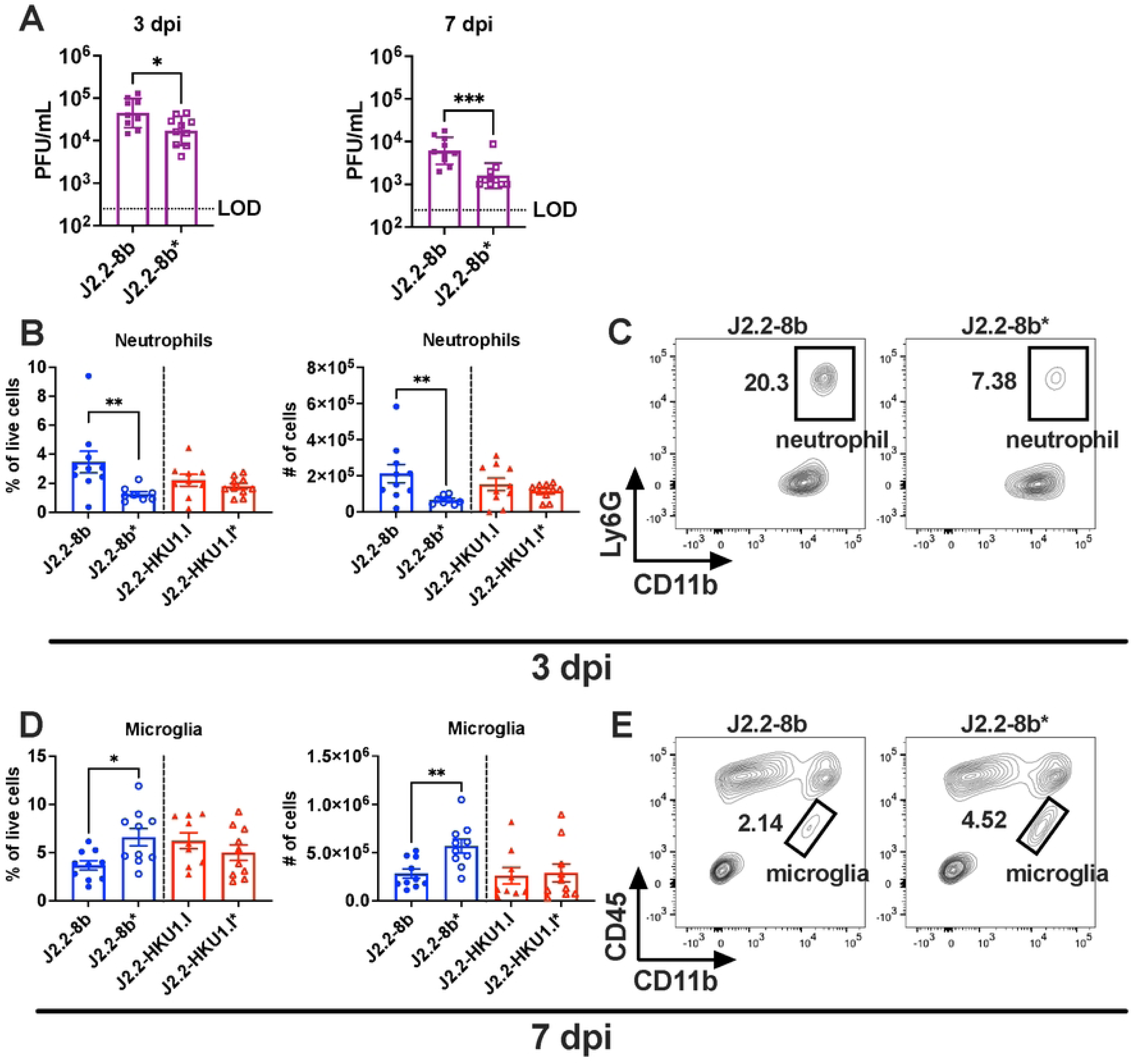
Changes in immune cell populations in J2.2-8b-infected brain. C57BL/6 mice were intracranially infected with 750 PFU of J2.2-8b or J2.2-8b* (A); J2.2-8b, J2.2-8b*, J2.2-HKU1.I or J2.2-HKU1.I* (B-E). The brains were harvested at 3 and 7 dpi for determination of infectious virus titers (A) and flow cytometric analysis of immune cell infiltration (B-E). Data in A are pooled from two independent experiments. Each point represents data obtained from an individual mouse. Data points are shown as geometric mean ± geometric SD. *P < 0.05, ***P < 0.001 by Student’s t test. (B, D) Frequency (left panel) and number (right panel) of neutrophils (B) and microglia (D) in infected brains are illustrated. Data are pooled from two independent experiments. Each point represents data obtained from an individual mouse. Data points are shown as mean ± SEM. *P < 0.05, **P < 0.01 by Student’s t test. (C, E) Representative flow plots of neutrophils (C) and microglia (E) in J2.2-8b-(left panel) or J2.2-8b*-infected (right panel) brains.

### The I proteins of hCoV-HKU1 and hCoV-OC43 exhibit different patterns of cellular localization

From our results, the hCoV-HKU1 but not the hCoV-OC43 I protein ameliorates J2.2 infection (**Figure 3 and 4**). To determine a mechanism for this difference in infection outcomes, we first assessed subcellular localization of the two I proteins. To test this, HeLa cells expressing murine carcinoembryonic antigen-related cell adhesion molecule (CEACAM1) (HeLa-MVR), the receptor for MHV were infected with J2.2-HKU1.I or J2.2-OC43.I. Infected HeLa-MVR were processed for immunofluorescence with an antibody against V5 tag to probe for the I proteins. The I protein of hCoV-HKU1 formed regions of strong fluorescence signals and punctate structures while diffused fluorescence signal across infected cells was observed for the I protein of hCoV-OC43 (**Figure 7**). Since the I proteins of MERS-CoV (protein 8b) and SARS-CoV-2 (protein 9b) were shown to interact with mitochondrial protein translocase of outer mitochondrial membrane 70 (TOM70) to varying extent [19,22,34], we stained for TOM70 in J2.2-HKU1.I- and J2.2-OC43.I-infected cells. As shown in **Figure 7A**, overlapping V5 and TOM70 fluorescence signals were observed in J2.2-HKU1.I-infected cells but not in cells infected with J2.2-OC43.I, suggesting that hCoV-HKU1 I protein may interact with TOM70. We further stained for Golgi matrix protein GM130 to determine if the I proteins of hCoV-HKU1 and hCoV-OC43 localize at the Golgi complex. We observed moderate overlapping between the V5 and GM130 signals in J2.2-HKU1.I-infected cells. However, co-localization between the I protein of hCoV-OC43 and GM130 was not detected (**Figure 7B**). Interestingly, condensed and perinuclear GM130 signal were observed in mock-infected cells while diffused GM130 signal was detected across J2.2-HKU1.I-infected cells with the loss of perinuclear localization, suggesting that the I protein of hCoV-HKU1 may disrupt the Golgi complex. Similarly, we identified diffused GM130 signal in some J2.2-OC43.I-infected cells albeit not as prominent as those infected with J2.2-HKU1.I. Overall, these data suggest that the I proteins of hCoV-HKU1 and hCoV-OC43 demonstrate distinct patterns of subcellular localization and possibly interact with different host proteins, potentially contributing to the unique infection outcomes in J2.2-HKU1.I- and J2.2-OC43.I-infected mice (**Figure 3**).

**Figure 7.**
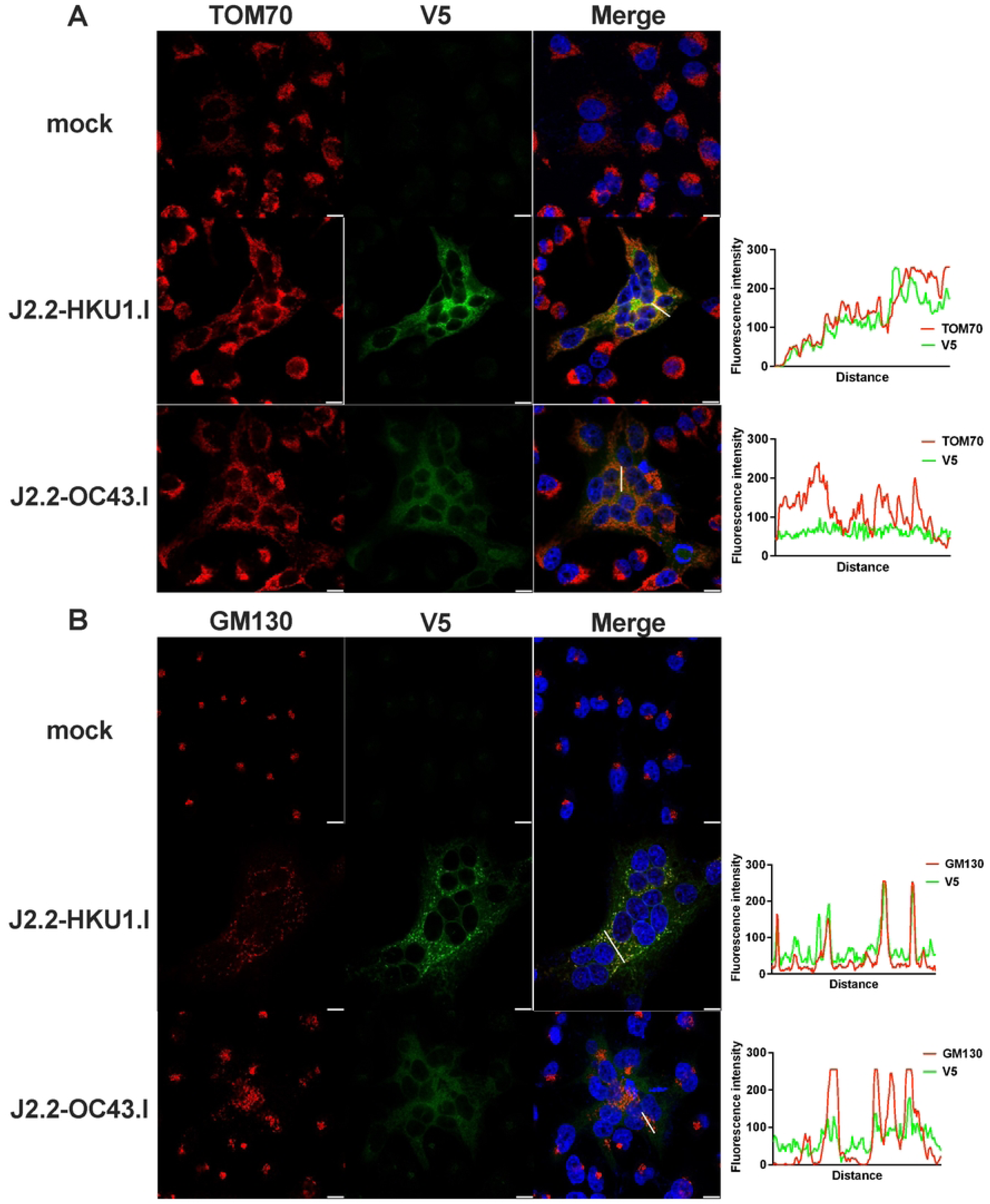
Distinct cellular localization of the I proteins of hCoV-HKU1 and hCoV-OC43. HeLa-MVR cells were infected with J2.2-HKU1.I or J2.2-OC43.I at an MOI of 0.01. Cells were washed and fixed at 16 hpi. The I protein signal was determined with an anti-V5 antibody (V5, green) while TOM70 (A) or GM130 (B) were stained in red. Intensity of fluorescence signals for the I protein (V5) and TOM70 (A) or GM130 (B) in infected cells along the indicated white line in the merged images are shown. Representative images from two independent experiments are shown.

## Discussion

Accessory proteins are the least conserved viral proteins among coronaviruses as compared to the structural and non-structural proteins, especially for viruses within different subgenera. However, betacoronaviruses of several subgenera encode the I protein as one of their accessory proteins, suggesting that the presence of the *I* gene and the expression of the I protein is a conserved feature among betacoronaviruses. Although the I protein is reported to act as an IFN antagonist, many CoVs encode multiple IFN antagonists with redundant functions [18,35–40]. Therefore, it is unlikely that the need for IFN antagonism alone selects for the expression of the I protein. Furthermore, the I gene is harbored within the N gene, which poses additional evolutionary constraints to the I protein, suggesting that undocumented I protein functions are important for CoV evolution. Interestingly, our previous study reported that the predicted structure of the I protein of MERS-CoV resembles the experimentally determined structure of the I protein of SARS-CoV-2 when interacting with its host protein TOM70 despite limited sequence homology [19]. This highlights the possibility that the I proteins of betacoronaviruses may form analogous structures and perform similar functions in the host during infection. Similarly, the I proteins of hCoV-HKU1 and hCoV-OC43 share low degree of sequence identity with the I proteins of MERS-CoV and SARS-CoV-2. Whether the I proteins of hCoV-HKU1 and hCoV-OC43 fold into structures comparable to that of the I proteins of MERS-CoV and SARS-CoV-2 requires further investigation. Although the I protein of hCoV-HKU1 showed signs of co-localization with TOM70 and the Golgi complex, these findings require validation with authentic hCoV-HKU1. The interaction between the I protein of hCoV-HKU1 and TOM70 or Golgi complex, if validated, should be further characterized to determine the mode of binding (direct vs. indirect, mediated by other host factors) and the molecular interactions responsible for the binding. Future research focusing on the structural perspectives of the I protein will elucidate critical virus-host interactions that are important for I protein-mediated pathogenesis.

Based on our findings, the I protein of hCoV-HKU1 causes reduction in virus replication *in vivo* (**Figure 3**) but not 17Cl-1 cells that induce minimal IFN after infection (**Figure 2A**), suggesting that the impact of the I protein of hCoV-HKU1 on virus replication is immune-mediated. In addition, IL-6, IFN and ISG expression were decreased in J2.2-HKU1.I-infected BMDM without significant reduction in viral RNA levels, suggesting that the hCoV-HKU1 I protein suppresses immune activation (**Figure 5**). Unlike J2.2-HKU.I, the I protein of MERS-CoV enhances J2.2 infection as it suppresses IFN and cytokine expression which leads to concomitant increase in virus replication [30]. While most of the characterized I proteins act as innate immune antagonists, the I protein of MHV-JHM has been shown to play a role in maintaining virion integrity and assembly in addition to immune antagonism [31], suggesting the pleiotropic nature of the I protein.

hCoV-HKU1 and hCoV-OC43 belong to the subgenus *Embecovirus*. These two viruses share highly homologous I protein sequences. Sequence alignment reveals that the I protein of hCoV-OC43 aligns to the C-terminus of the hCoV-HKU1 I protein (**Figure S1**), suggesting that the difference in the phenotype between J2.2-HKU1.I- and J2.2-OC43.I-infected mice maybe attributed to the N-terminus of the hCoV-HKU1 I protein. A previous study demonstrated that the N-terminus of the I protein of MERS-CoV is critical for IFN antagonism and enhanced disease after J2.2 infection [30], hinting a role of the N-terminal domain in mediating I protein function. This earlier study also reported the presence of multiple epitope-tagged protein products expressed by the *I* gene of MERS-CoV in the context of J2.2 infection [30]. We did not detect multiple protein bands in J2.2-8b-infected cells in this study due to the use of an antibody that targets the N-terminus of protein 8b [18]. Similarly, we detected multiple protein products in J2.2-HKU1.I-infected cells when an antibody targeting the C-terminal epitope tag was used (**Figure 1C**). The faster-migrating bands likely represent protein products in which translation initiates at alternative start codons downstream of the first start codon (**Figure S1**). In particular, a minor protein product of small molecular weight was present in J2.2-HKU1.I*-infected cells (**Figure 1C**). Based on these observations, we conclude that the phenotypic differences observed between J2.2-HKU1.I and J2.2-HKU.1* are likely contributed by the full-length I protein.

MHV-J2.2 is a neurotropic strain of MHV that causes infection of the central nervous system (CNS). This study investigates the role of I proteins of hCoV-HKU1 and hCoV-OC43 in pathogenesis, both of which are respiratory hCoVs. The mismatch in the site of infection (CNS vs airway) and tissue-specific immune responses may not reflect the phenotype of the I proteins in natural (airway) infections. Furthermore, protein 8b of MERS-CoV is reported to play opposite roles in pathogenesis in the context of MERS-CoV and J2.2 infection. We previously showed that the presence of protein 8b prevents lethal disease after MERS-CoV infection but contributes to enhanced disease in the context of J2.2 infection [19]. In fact, the role of protein 8b in exacerbating J2.2 infection has been independently reported in a separate study [30], suggesting that the I protein of hCoVs may contribute to pathogenesis in a virus-specific context. Combining with this study, future investigation in the context of authentic hCoV-HKU1 infection would offer critical insights in the virus- and/or tissue-specific role of the I protein. The use of authentic hCoV-HKU1 remains a technical challenge as it is reported to only replicate in primary human airway cells [41], rendering it challenging for propagation and genetic manipulation.

In conclusion, we compared the I proteins of three hCoVs in relation to pathogenesis in the context of J2.2 infection. We characterized the impacts of the different I proteins on the virological properties of MHV-J2.2 and the host immunological changes in experimentally infected animals. These results further affirm the unique roles of the I protein of various CoVs in pathogenesis and reinforce the notion that accessory proteins are important in modulating CoV life cycle in a virus-specific fashion.

## Materials and Methods

### Mice, cells and viruses

Specific-pathogen-free C57BL/6 mice were purchased from Charles River Laboratories and maintained in a specific pathogen-free facility at Rutgers University under standard conditions of dark/light cycle, ambient temperature, and humidity. Five- to seven-week-old male mice were used in all experiments. Female mice exhibit similar clinical outcomes after JHMV infection, but results are less consistent. All animal studies were approved by the Institutional Animal Care and Use Committee (IACUC) of Rutgers University and follow guidelines of the *Guide for the Care and Use of Laboratory Animals*. 17Cl-1 mouse fibroblasts, HeLa cells expressing the MHV receptor carcinoembryonic antigen-related cell adhesion molecule 1 (HeLa-MVR) and BHK-21 cells were grown in Dulbecco’s modified Eagle medium (DMEM) supplemented with 10% fetal bovine serum (FBS), 100 U/mL penicillin and streptomycin, L-glutamine, sodium pyruvate, HEPES, and non-essential amino acids and maintained at 37°C. BMDMs were harvested from C57BL/6 mice and differentiated in DMEM media supplemented with 10% FBS, M-CSF (20 ng/mL), 100 U/mL penicillin and streptomycin, L-glutamine, and sodium pyruvate. BMDM media were changed every other day following the fourth day of differentiation and infected at 7 or 8 days post-differentiation. A recombinant version of the neuroattenuated J2.2-V-1 variant of JHMV (MHV-J2.2) was propagated in 17Cl-1 cells. Virus titers of MHV-J2.2 were determined in HeLa-MVR cells. Studies involving the use of recombinant viruses have been approved by the Institutional Biosafety Committee of Rutgers University.

### Mouse infection

Five- to seven-week-old male mice were anesthetized with ketamine-xylazine and inoculated intracranially with 750 PFU of MHV in 30 µL of DMEM. Mice were monitored and weighed daily following inoculation. Clinical scoring for MHV infection was based on the following criteria: 0, asymptomatic; 1, limp tail, mild hunching; 2, wobbly gait with mild righting difficulty, hunching; 3, hind-limb paresis and extreme righting difficulty; 4, hind-limb paralysis; and 5, moribund.

### Infection of cell and virus plaque assay

Cells were washed with PBS once before infection. Viruses were diluted in DMEM for infection at the indicated MOI. Cells were incubated with virus inoculum for 1 h at the conditions described below. After infection, virus inoculum was removed and DMEM supplemented with 10% FBS was added to the well and incubated until time of harvest. For plaque assay, cells were frozen and mice were euthanized and transcardially perfused with PBS at the indicated times. Organs were harvested and homogenized, while cells were thawed prior to clarification by centrifugation and titering. Virus or tissue homogenate were serially diluted in DMEM. Twelve-well plates of HeLa-MVR were inoculated with at 37°C in 5% CO_2_ for 1 h and gently rocked every 15 min. After 1 h of incubation, virus inoculum was removed and DMEM supplemented with 10% FBS was added to the well and incubated for 16 h. After 16 h, the supernatant was removed and plates were overlaid with 0.6% agarose containing 2% FBS and 0.1% neutral red and incubated for 4 h. Plaques were counted without further treatment after 4 h of incubation. Viral titers were quantified as PFU/mL.

### Generation of mutant viruses

OC43.I, OC43.I*, HKU1.I or HKU1.I* sequences were introduced into J2.2 BAC at the ORF4 region by a two-step linear lambda red recombination process [42–44]. The first step removed and replaced the ORF4 sequence with GalK-Kan selection marker while the second step removed and replaced the GalK-Kan selection marker with the desired insert sequences prepared by PCR amplification followed by purification. In brief, GalK-Kan selection marker was PCR amplified from pYD-C225 [44] and gel purified. Gel-purified GalK-Kan fragments were transformed into SW102 cells carrying the J2.2 BAC by electroporation, for linear lambda red recombination. Successful recombinants were selected on Kanamycin resistance culture plates. Verified recombinants carrying GalK-Kan cassette were further introduced with the corresponding inserts by electroporation for a second round of linear lambda red recombination. Successful recombinants were selected using 2-deoxy-galactose-based culture plates, and sequence identity was verified by BAC sequencing. GalK-Kan selection markers were amplified with the following primers: forward, 5’-CTCTCCTGGAAAGACAGAAAATCTAAACAATTTATAGCATTCTCATTGCTACTTTGCTCCTCTAGAGGGCAGCAAGTAGTT**cctgttgacaattaatcatcg**-3′; reverse, 5′-TACTTCGGCAAGTGCCTAAGTGTGTATGGACGGCCAGAATTAAGATGAGGTTTAGAACTAGTAATATAATCTAGAGT**ctcagcaaaagttcgattta**-3′. Lightface capital letters represent sequence flanking the area of interest in the J2.2 BAC; sequences complementary to pYD-C225 are shown as boldface lowercase letters.

Primers amplifying HKU1.I and HKU1.I* were as follows: forward, 5′-CTCTCCTGGAAAGACAGAAAATCTAAACAATTTATAGCATTCTCATTGCTACTTTGCTCCTCTAGAGGGCAGCAAGTAGTTatgctggaagtagaagctcctctgg-3′; reverse, 5′-AGGTTTAGAACTAGTAATATAATCTAGAGT**TTACGTAGAATCGAGACCGAGGAGAGGGTTAGGGATAGGCTTACC**caccagaggtaggggttctattgcc -3′. Primers amplifying OC43.I and OC43.I* were as follows: forward, 5′-TGGAAAGACAGAAAATCTAAACAATTTATAGCATTCTCATTGCTACTTTGCTCCTCTAGAGGGCAGCAAGTAGTTatggcaaccagcgtcaactgctgcc-3′; reverse, 5′-AGGTTTAGAACTAGTAATATAATCTAGAGT**TTACGTAGAATCGAGACCGAGGAGAGGGTTAGGGATAGGCTTACC**caccagaggtaggggttctattgcc-3′. The capital letters represent sequence of the area where recombination takes place (flanking ORF4 for MHV) while the boldface capital letters represent the V5 sequence and the lowercase letters represent sequence complementary to the corresponding I protein sequences for PCR amplification with synthesized plasmids as templates.

### Recovery of recombinant viruses and virus propagation

Two micrograms of the indicated J2.2 BACs was transfected into BHK-21 cells with Lipofectamine 3000 (Invitrogen) in a six-well plate according to the manufacturer’s protocol. Cells were monitored daily for cytopathic effects (CPE). Cultures were harvested when CPE was >50% by freezing at −80°C. MHV-J2.2 were further passaged in 17Cl-1 cells in DMEM supplemented with 10% FBS. Virus titer was determined by plaque assay as described in the previous section.

### Protein extraction and western blot analysis

17Cl-1 cells infected with J2.2 were lysed with RIPA buffer at the indicated time points and mixed with gel-loading dye for 10 min at 95°C before electrophoresis. Cell lysates were resolved by SDS-PAGE. Specific proteins were probed with the following primary antibodies: rabbit α-ORF8b polyclonal antibody [18], mouse α-N monoclonal antibody (Cat.# MABF2751; Millipore Sigma), mouse α-V5 monoclonal antibody (Cat.# R960-25; Invitrogen) or mouse α-actin monoclonal antibody (Cat.# MA1-140; Thermo Fisher Scientific). Bands were visualized with α-mouse (Cat. # 405306; BioLegend) or α-rabbit (Cat. # 406401; BioLegend) IgG secondary antibody conjugated with horseradish peroxidase (HRP) followed by substrate incubation (Cat. # 32106; Thermo Fisher Scientific).

### RNA isolation and qRT-PCR

Cells and tissues were homogenized in TRIzol (Invitrogen) for RNA isolation as specified by the manufacturer’s protocol. Isolated RNA was subject to DNase I treatment (Invitrogen) and reverse transcribed using the SuperScript IV First-Strand Synthesis System (Invitrogen). mRNA levels were determined after normalizing with HPRT by the ΔCt method. Specific primer sets used for qPCR were previously described [45,46]. Viral genomic RNA was detected with the following primers: forward, 5’-AGGGAGTTTGACCTTGTTCAG-3’; reverse, 5’-ATAATGCACCTGTCATCCTCG-3’

### RNA sequencing and analysis

RNA libraries were prepared using the NEB Next Ultra II RNA Library Prep Kit for Illumina (Catalog #E7770) with Poly(A) selection. In brief, 100 ng of total RNA in 50 μL of nuclease-free water was denatured at 65°C for 5 minutes, rapidly cooled to 4°C, and incubated at room temperature for 5 minutes with Oligo(dT) beads to capture mRNA. The beads were washed, and the bound mRNA was eluted in tris buffer. A second round of Oligo(dT) bead purification was performed, with final elution in first-strand synthesis buffer.

cDNA synthesis was carried out in two stages: first-strand synthesis, followed by second-strand synthesis, with purification using AMPure XP beads (Beckman Coulter, Cat. # A63881). The resulting cDNA underwent end-repair, adaptor ligation, and barcoding with unique i7 and i5 index primers. A subsequent round of AMPure XP bead purification was performed to refine the libraries. Library quality was assessed using the Agilent TapeStation system (Part #G2992AA) and quantified with a Qubit fluorometer. The libraries were then pooled in equimolar ratios and sequenced on the NovaSeq X Plus platform using 1.5B, 200 cycle kit with paired-end configuration, generating an average of 50 million reads per sample.

Raw transcriptome reads were assessed for quality control (FASTQC v0.11.9) and trimmed for quality and adapter contaminants (cutadapt v 3.4). Trimmed reads were aligned to the Mus musculus genome (GRCm38) using STAR (v2.7.10a), followed by transcript abundance calculation and hit count extraction with StringTie (v2.2.1) and featureCounts (v2.0.1) respectively. Hit count normalization and differential gene expression group cross-comparisons were performed using DESeq2 (v1.44.0). Significant differentially expressed gene thresholds were set at FDR adjusted p <0.05. Qiagen IPA (v01-23-01) Expression Analysis based on experimental log ratio and a p-adjusted cutoff of < 0.05 was used for pathway analysis. Raw RNA sequencing data were deposited to NIH Gene Expression Omnibus (Project ID: GSE291309).

### Flow cytometric analysis

Animals were anesthetized with ketamine-xylazine and perfused transcardially with 10 mL PBS. Brains were removed, minced, and digested in HBSS buffer consisting of 2% fetal calf serum, 25 mM HEPES, 1 mg/mL collagenase D (Roche), and 0.1 mg/mL DNase (Roche) at 37°C for 30 min. Single-cell suspensions were prepared by passage through a 70 µM cell strainer. Cells were enumerated with Countess 3 (Invitrogen). Cells were then washed and blocked with 1 µg α-CD16/α-CD32 (2.4G2) antibody at 4°C for 20 min and surface stained with the following antibodies at 4°C for 30 min: V450 α-CD45 (clone 30-F11; BioLegend); BV605 α-CD11b (clone M1/70; BioLegend); PE α-Ly6G (clone 1A8; BioLegend); BV785 α-Ly6C (clone HK1.4; BioLegend) and LIVE/DEAD Fixable Blue Dead Cell Stain Kit (Thermo Fisher Scientific). All flow cytometry data were acquired using a BD FACS Symphony and analyzed with FlowJo software.

### Immunofluorescence

HeLa-MVR cells seeded on cover slips were mock infected or infected with J2.2-HKU1.I or J2.2-OC43.I at an MOI of 0.01. At 16h post infection, cells were washed with PBS and fixed with 4% paraformaldehyde for 20 min at room temperature. Cells were washed three times with PBS and permeabilized in 0.75% of Triton X-100 for 20 min. Permeabilized cells were blocked with 1% BSA for 1 h at room temperature and stained with primary antibodies (α-V5, 1:120 dilution, Invitrogen Cat. # R960-25; α-TOM70, 1:120 dilution, Abcam Cat. # ab289977) at 4 degrees overnight. Cells were washed three times with PBS the next day and stained with secondary antibodies at room temperature for 1 h. Cells on cover slips were washed three times with PBS and mounted with Vectashield (Cat. # H-1800) on slides for imaging with Leica STED8 confocal microscope.

### Statistical analysis

Student’s *t*-test was used to analyze differences in mean values between groups. All results are expressed as mean ± SEM except for virus titers and viral RNA levels where data are represented as geometric mean ± geometric SD. *P*-values of <0.05 were considered statistically significant. \**P* < 0.05, \*\**P* < 0.01, and \*\*\**P* < 0.001. Differences in mortality were analyzed using log-rank (Mantel-Cox) survival tests.

## Acknowledgements

We thank Shea Lowery for reviewing the manuscript. This study is supported in part by grants from the NIH (R00 AI170996, L.-Y.R.W.). RNA sequencing was performed at the Molecular and Genomics Informatics Core Facility at New Jersey Medical School.

C.D.G, R.R., A.S. and L.-Y.R.W. conceived the work and designed the experiments; C.D.G., I.P., S.C., Y.P.-T. and A.O. acquired the data; A.M., A.S. and L.-Y.R.W. analyzed the data; L.-Y.R.W. wrote the manuscript with contribution from all authors.

## Supplemental information

**Figure S1.**
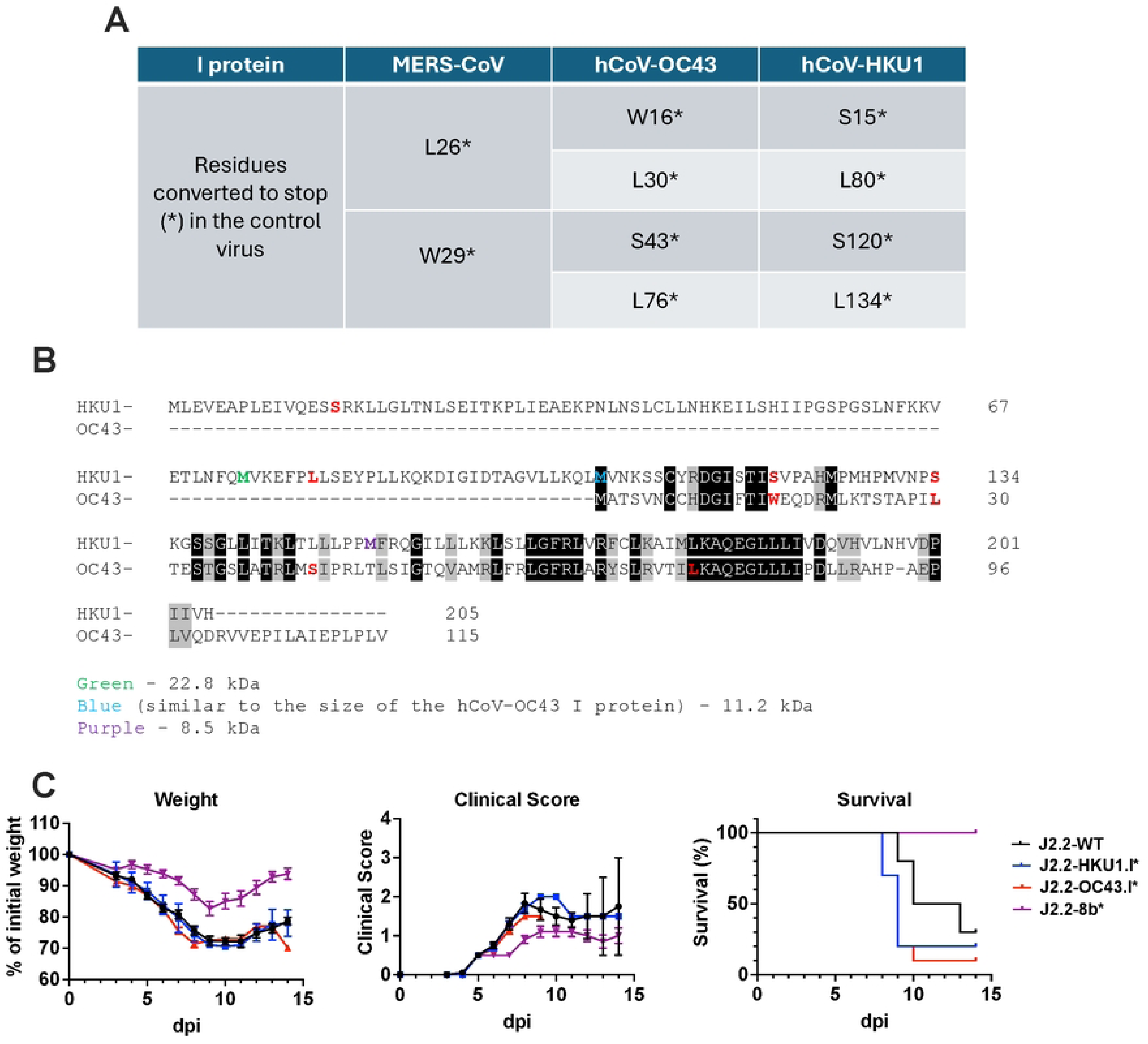
I protein and control sequences used in this study. (A) Table showing the premature stop mutations introduced to the control viruses. (B) Sequence alignment between the I proteins of hCoV-HKU1 and hCOV-OC43. Sequences were derived from GenBank (accession number AY597011 and AY391777 for the I proteins of hCoV-HKU1 and hCoV-OC43, respectively). Methionine residues highlighted in green, blue and purple in the sequence of the hCoV-HKU1 I protein represent the putative start of the protein products indicated by the arrowheads of the corresponding color shown in Figure 1C. Putative starts are predicted based on the molecular weight of the protein products. Residues converted to stop codons in J2.2-OC43.I* and J2.2-HKU1.I* are highlighted in red. Shades of black and gray represent identical and similar amino acid residues between the I protein of hCoV-HKU1 and hCoV-OC43, respectively. (C) Percent of initial weight (left panel), clinical score (middle panel) and survival (right panel) of C57BL/6 mice intracranially infected with 750 PFU of J2.2, J2.2-HKU1.I*, J2.2-OC43.I* or J2.2-8b*. Data are pooled from two independent experiments (n = 10 for each group). Data points are shown as mean ± SEM for the weight curve and the panel for clinical score.

**Figure S2.**
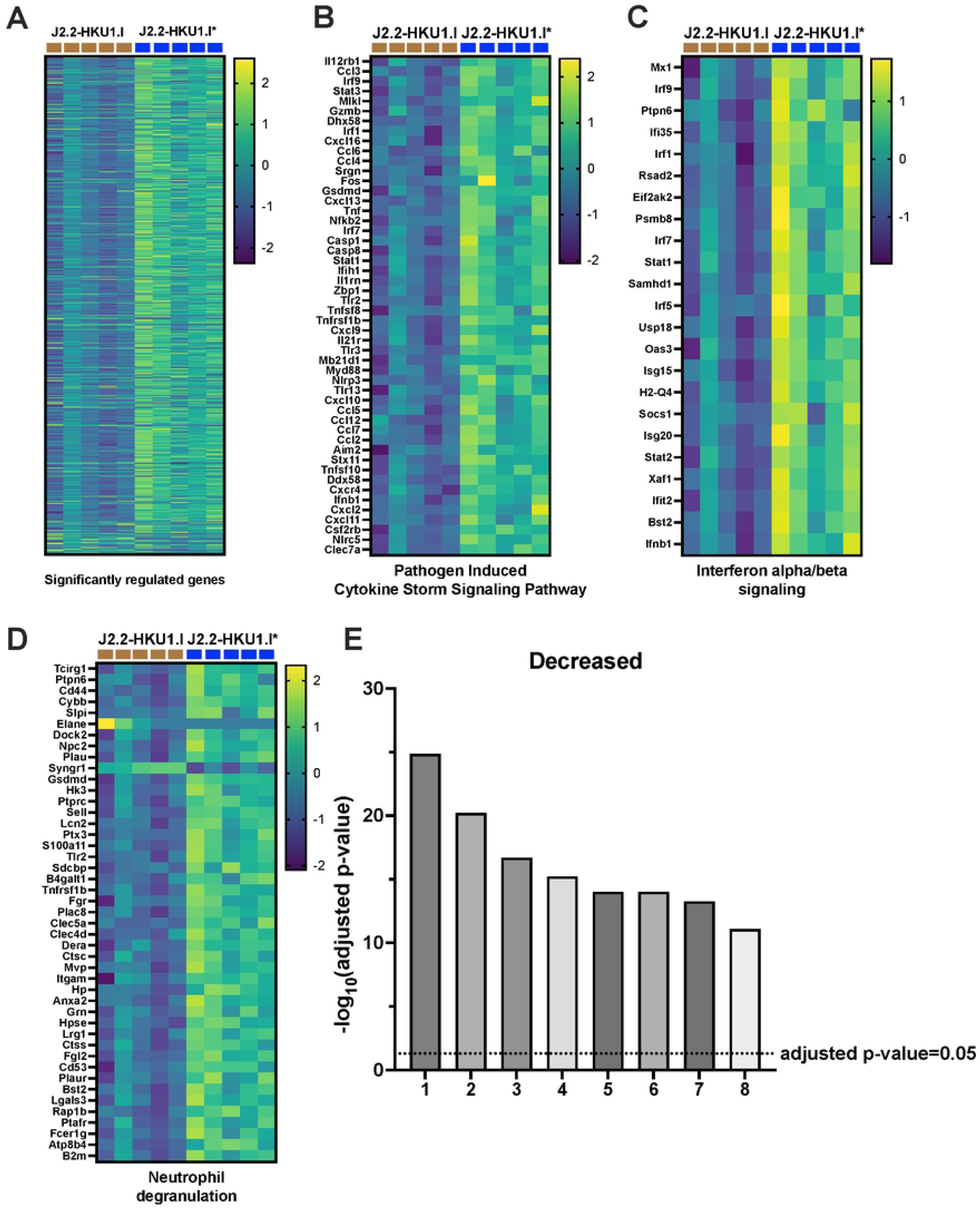
Brains infected with J2.2-HKU1.I show downregulation of pathways related to cytokine signaling and antiviral response. C57BL/6 mice were intracranially infected with 750 PFU of J2.2-HKU1.I or J2.2-HKU1.I*. The brains were harvested at 3 dpi for isolation of total RNA followed by RNA sequencing. Genes with adjusted P<0.05 were selected for further analysis. Heat maps plotting log_10_ z-score of all significantly regulated genes (A), genes involved in pathogen induced cytokine signaling pathway (B), interferon alpha/beta signaling (C) and neutrophil degranulation (D) at 3 dpi. (E) Ingenuity Pathway Analysis (Qiagen) was used to analyze altered biological pathways (J2.2-HKU1.I vs. J2.2-HKU1.I*) at 3 dpi. Pathways with adjusted *P* < 0.05 are considered significant. 1: Pathogen Induced Cytokine Storm Signaling Pathway; 2: Interferon alpha/beta signaling; 3: Neutrophil degranulation; 4: ISGylation Signaling Pathway; 5: Interferon gamma signaling; 6: Role of PKR in Interferon Induction and Antiviral Response; 7: IL-27 Signaling Pathway; 8: Macrophage Classical Activation Signaling Pathway

**Figure S3.**
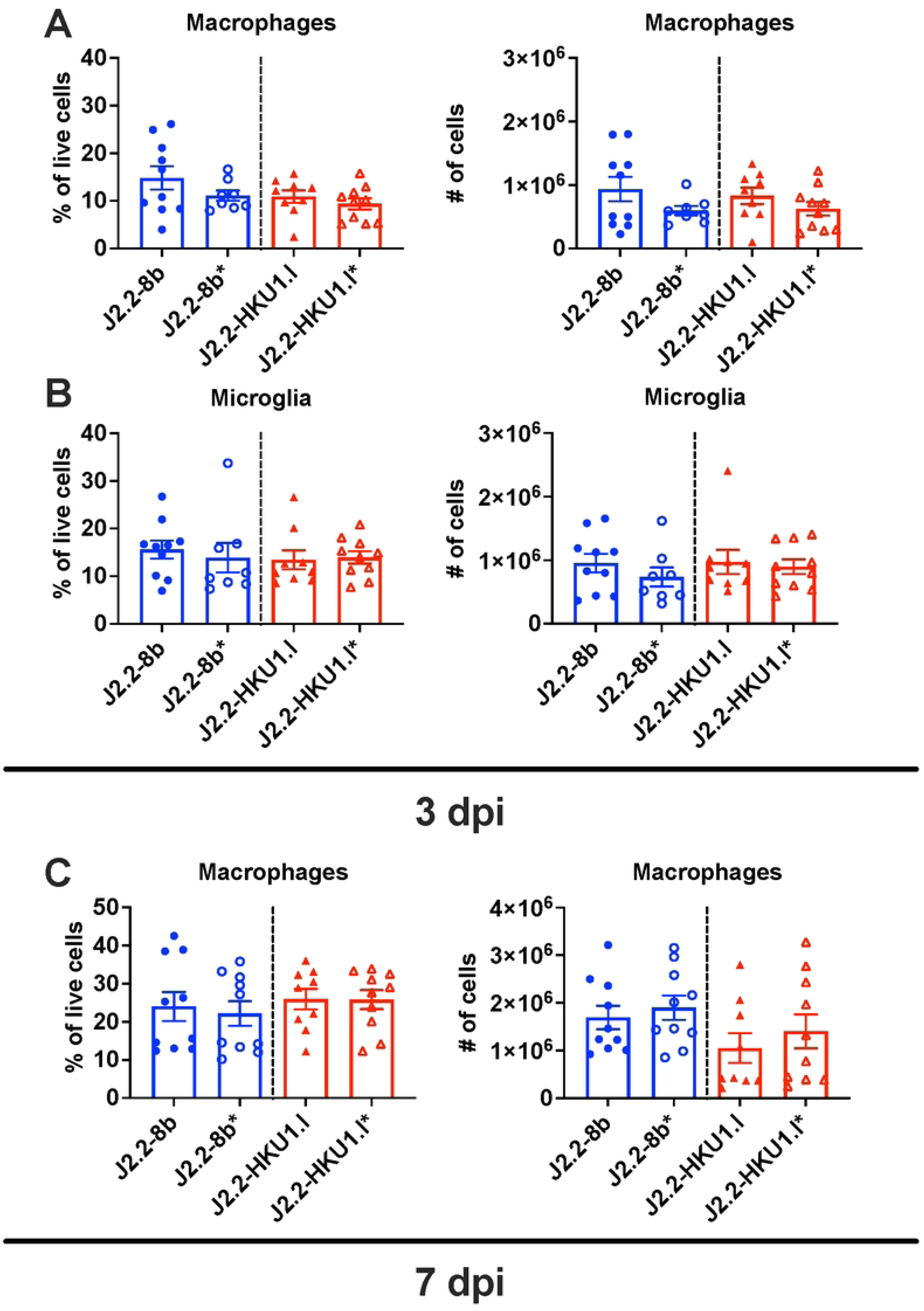
Immune cell profiling in the brains infected with J2.2 expressing the I proteins of MERS-CoV or hCoV-HKU1. C57BL/6 mice were intracranially infected with 750 PFU of J2.2-8b, J2.2-8b*, J2.2-HKU1.I or J2.2-HKU1.I*. The brains were harvested at 3 and 7 dpi for flow cytometric analysis of immune cell infiltration. Frequency (left panel) and number (right panel) of macrophages (A), microglia (B) at 3 dpi and macrophages at 7 dpi in infected brains are illustrated. Data are pooled from two independent experiments. Each point represents data obtained from an individual mouse. Data points are shown as mean ± geometric SEM.

